# rphenoscate: An R package for semantic-aware evolutionary analyses of anatomical traits

**DOI:** 10.1101/2023.02.19.528613

**Authors:** Diego S. Porto, Sergei Tarasov, Caleb Charpentier, Hilmar Lapp, James P. Balhoff, Todd J. Vision, Wasila M. Dahdul, Paula M. Mabee, Josef Uyeda

## Abstract

1. Organismal anatomy is a complex hierarchical system of interconnected anatomical entities often producing dependencies among multiple morphological characters. Ontologies provide a formalized and computable framework for representing and incorporating prior biological knowledge about anatomical dependencies in models of trait evolution. Further, ontologies offer new opportunities for assembling and working with semantic representations of morphological data.
2. In this work we present a new R package—*rphenoscate*—that enables incorporating ontological knowledge in evolutionary analyses and exploring semantic patterns of morphological data. In conjunction with *rphenoscape* it also allows for assembling synthetic phylogenetic character matrices from semantic phenotypes of morphological data. We showcase the new package functionalities with three data sets from bees and fishes.
3. We demonstrate that ontology knowledge can be employed to automatically set up ontology-informed evolutionary models that account for trait dependencies in the context of stochastic character mapping. We also demonstrate how ontology annotations can be explored to interrogate patterns of morphological evolution. Finally, we demonstrate that synthetic character matrices assembled from semantic phenotypes retain most of the phylogenetic information of the original data set.
4. Ontologies will become an increasingly important tool not only for enabling prior anatomical knowledge to be integrated into phylogenetic methods but also to make morphological data FAIR compliant—a critical component of the ongoing ‘phenomics’ revolution. Our new package offers key advancements toward this goal.

## 1 Introduction

Biological realism in models of trait evolution—*i*.*e*., accurate modeling of biological processes underlying trait changes through time—is often an overlooked but important feature in phylogenetic comparative modeling (Boyko and Beaulieu, 2021). For example, it is common in statistical phylogenetics to treat each character as an independent realization of the evolutionary process. While this assumption may be questionable for molecular data, it is certainly dubious for morphological data. Nevertheless, this assumption is commonly applied in morphological analyses (see discussions in Lewis, 2001; Wright, 2019). Non-independence among anatomical traits can result from multiple causes (*e*.*g*., see the distinction among *biological, semantic* and *ontological* dependencies in Vogt, 2018a) and alternative models have been proposed to properly deal with them (*e*.*g*., Tarasov, 2019, 2022). While researchers often attempt to at least partially deal with such challenges via expert character construction, there is a pressing need for such knowledge to be repeatable and computable. What if we could reliably inform phylogenetic models with prior knowledge on anatomical trait relationships, including potential biological and/or logical dependencies, in a repeatable and computable framework? In this paper, we present a new R package for addressing this challenge, *rphenoscate*, that enables semantically-aware evolutionary analyses by integrating morphological knowledge present in anatomy ontologies.

The ‘dependency problem’—how to code and model dependent traits—often associated with missing or inapplicable characters, is a longstanding issue in phylogenetics with morphological data (the ‘tail color problem’ from Maddison, 1993 as referred in Tarasov, 2019) and has received considerable attention in recent years (Tarasov, 2019, 2022; Goloboff et al., 2021; Hopkins and St. John, 2021; Simões et al., 2022). This issue is especially relevant if we want to improve the biological realism of evolutionary models for morphological traits, as organismal anatomy is highly structured and phylogenetic characters often refer to multiple anatomical entities and/or phenotypes exhibiting complex hierarchical relations (Porto et al., 2021, 2022). Advances in model-based phylogenetics now allow researchers to employ different models and coding strategies to deal with character dependencies (Tarasov, 2019, 2022). Although there still is a discussion on how to properly set up these models and represent dependencies in a coding scheme (see Goloboff et al., 2021; Simões et al., 2022), ontologies can offer an answer to ‘what the dependencies are’. Thus, anatomy ontologies are important sources of computable biological knowledge about organismal anatomy and are the key to enabling reproducibility and integration of biological knowledge into phylogenetic workflows.

Ontologies are formal representations of domain knowledge using structured vocabularies (Balhoff et al., 2010; Dahdul et al., 2010b, 2012; Vogt, 2018a,b). Anatomy ontologies, in particular, allow one to express knowledge about different anatomical concepts in a particular group of organisms (Dahdul et al.,

2010b). For example, ontologies can formalize that the anatomical concept ‘dorsal fin ray’ is *part of* ‘dorsal fin’. Therefore, the condition of a character representing a ‘dorsal fin ray’ (*e*.*g*., shape or number of rays) depends on the presence of a ‘dorsal fin’. Despite being a rather simple statement for a trained fish anatomist, this type of biological knowledge is crucial for computers to be able to autonomously reason about trait evolution—yet such relationships are increasingly likely to be lost as analyses transition from expertly curated data sets to large automated data syntheses. If trait dependencies are not accounted for, for example, this can result in overestimating the true amount of evolutionary change, potentially affecting divergence time estimates using fossilized birth-death models (Ronquist et al., 2012; Wright et al., 2022). Additionally, ignoring dependencies can result in biologically unrealistic combinations of states at internal nodes when performing ancestral character state reconstruction with multiple traits (Forey and Kitching, 2000; Tarasov, 2019; Boyko and Beaulieu, 2021). Even when these are not the direct target of inference, many comparative methods such as state-dependent diversification models (FitzJohn, 2012) and character correlation tests (e.g. Pagel, 1994) integrate over these ancestral probabilities and therefore can be affected by considering implausible character histories. Therefore, employing appropriate models is not only desirable for improving biological realism but also necessary to avoid misleading results. By providing tools that automate model specification when dependent traits are present using the formalized knowledge in anatomy ontologies (*e*.*g*., Tarasov, 2019, 2022), our new R package enables researchers to quickly and easily structure biologically-plausible models of character evolution for phenomic-scale matrices.

Besides informing models, ontologies open up new questions for researchers interested in the evolution of morphological traits. Dependencies among anatomical entities—and the phylogenetic characters proposed from them—can be of several types (Vogt, 2018a; see some useful definitions of concepts discussed along the text in Table 1). Using ontology annotations to phylogenetic characters one can, for example, automatically assemble all characters representing traits that are *part of* ‘cranium’ (*e*.*g*., bones: ‘endopterygoid’, ‘parasphenoid’, ‘parietal’), *is a* type of ‘anatomical projection’, or *develops from* the ‘mesoderm’ in a fish, and then test to see if different bones from the same cluster evolve at similar rates. Alternatively, one can use such clusters to further investigate if the phylogenetic characters linked to the anatomical entities share other parameters in their evolutionary models (*e*.*g*., transition bias). For example, are certain types (*is a*) of anatomical entities or entities belonging to a certain body region (*part of*) more prone to be lost during evolution? These are just a few examples of the utility of ontologies in evolutionary analyses (see also Dahdul et al., 2010a; Ramírez and Michalik, 2014; Vogt, 2018a,b; Tarasov et al., 2019; Tarasov, 2019; Porto et al., 2022).

**Table 1:**
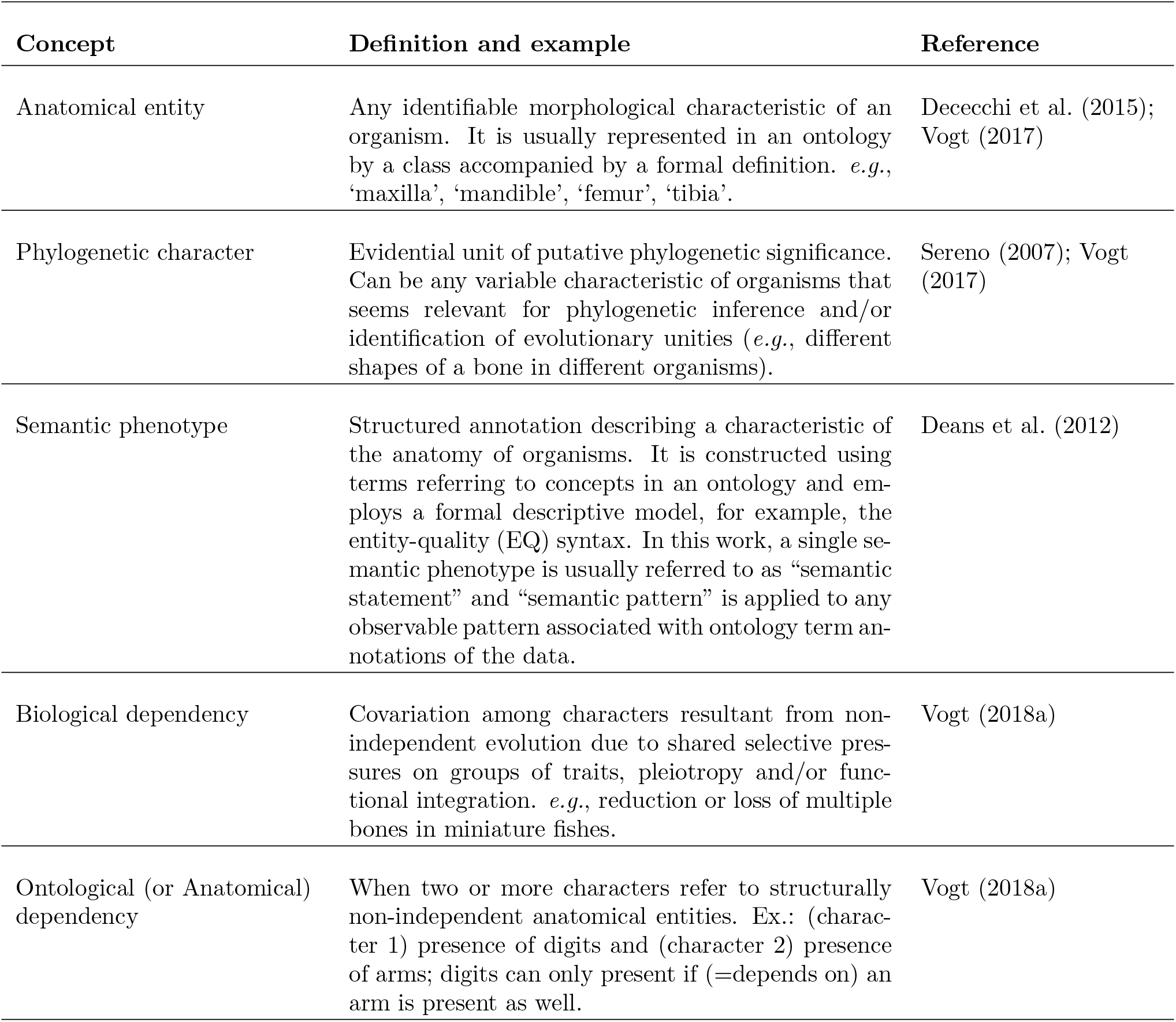
Glossary of some important concepts and their definitions.

| Concept | Definition and example | Reference |
| --- | --- | --- |
| Anatomical entity | Any identifiable morphological characteristic of an organism. It is usually represented in an ontology by a class accompanied by a formal definition. *e.g.*, ‘maxilla’, ‘mandible’, ‘femur’, ‘tibia’. | Dececchi et al. (2015); Vogt (2017) |
| Phylogenetic character | Evidential unit of putative phylogenetic significance. Can be any variable characteristic of organisms that seems relevant for phylogenetic inference and/or identification of evolutionary unities (*e.g.*, different shapes of a bone in different organisms). | Sereno (2007); Vogt (2017) |
| Semantic phenotype | Structured annotation describing a characteristic of the anatomy of organisms. It is constructed using terms referring to concepts in an ontology and employs a formal descriptive model, for example, the entity-quality (EQ) syntax. In this work, a single semantic phenotype is usually referred to as “semantic statement” and “semantic pattern” is applied to any observable pattern associated with ontology term annotations of the data. | Deans et al. (2012) |
| Biological dependency | Covariation among characters resultant from non-independent evolution due to shared selective pressures on groups of traits, pleiotropy and/or functional integration. *e.g.*, reduction or loss of multiple bones in miniature fishes. | Vogt (2018a) |
| Ontological (or Anatomical) dependency | When two or more characters refer to structurally non-independent anatomical entities. Ex.: (character 1) presence of digits and (character 2) presence of arms; digits can only present if (=depends on) an arm is present as well. | Vogt (2018a) |

Furthermore, ontologies not only offer a solution for the longstanding ‘dependency problem’ among anatomical traits in phylogenetics (Tarasov, 2019) but also, an interoperable framework for representing morphological knowledge and integrating it with other knowledge types. Based on theoretical and practical grounds, recent works have suggested new schema for employing morphological data in phylogenetics (Vogt, 2018a,b), for example, by using semantically-enriched character matrices (*e*.*g*., Ramírez et al., 2007; Stefen et al., 2022), semantic instance anatomies (*e*.*g*., Vogt, 2018a,b, 2019), or semantic phenotypes (*e*.*g*., Deans et al., 2012; Balhoff et al., 2010, 2014). By representing organismal anatomy in a semantically-aware format (*i*.*e*., ontology-annotated) and moving beyond the standard phylogenetic character matrices, it is possible to make morphological data more easily reusable, parsable, and integrated across different studies and domains. Some new uses include, but are not restricted to, building synthetic character matrices from multiple sources (Dececchi et al., 2015; Jackson et al., 2018; Eliason et al., 2019), inferring candidate genes for novel phylogenetic traits (Edmunds et al., 2016), and graph-based phylogenetic algorithms (Vogt, 2018a,b). To enable such analyses, the Phenoscape project (https://kb.phenoscape.org/) has developed key demonstrations of the use of ontologies in the development of a logical model of homology (Mabee et al., 2020) and inference of candidate genes from phylogenetic traits (Edmunds et al., 2016; Manda et al., 2015), as well as gold standards for curation (Dahdul et al., 2018). These grew from the development of one of the first multispecies anatomy ontologies for the biodiversity sciences. Their initial teleost fish ontology (Dahdul et al., 2010b) grew into a vertebrate ontology (Dahdul et al., 2012) and merged into the Uberon anatomy ontology (Haendel et al., 2014), used herein. As part of these demonstrations, they developed an expert-curated database of semantic phenotypes (*i*.*e*., the Phenoscape Knowledgebase, *e*.*g*., Manda et al., 2015) for more than 4,800 extant and extinct teleost fishes. Our new R package capitalizes on this knowledgebase to provide some tools for exploring new phylogenetic applications of semantically-aware anatomical data.

In this study, we implemented several tools for performing semantic-aware evolutionary analyses and exploring semantically-aware morphological data in a new R package *rphenoscate*. These tools include functions for automatically setting up evolutionary models for dependent traits based on a reference anatomy ontology, a phylogenetic data set, and character annotations to ontology terms. We integrated the new package with previous R packages tailored to work with phylogenetic data and ontologies, such as *rphenoscape* (https://github.com/phenoscape/rphenoscape), *ontologyIndex* (Greene et al., 2017), *ontoFAST* (Tarasov et al., 2022), and the PARAMO pipeline (Tarasov et al., 2019). We provide tools to prepare data and models for evolutionary analyses (*e*.*g*., stochastic character mapping) that can be performed in R (*e*.*g*., corHMM, Boyko and Beaulieu, 2021) or in RevBayes (Hühna et al., 2016). *rphenoscate* also offers functions for importing and visualizing results, including tools for investigating relationships among anatomy ontology term annotations. *rphenoscate* further offers tools for assembling synthetic character matrices from semantic phenotypes available from the Phenoscape Knowledgebase (Phenoscape KB). We showcase the package functionality with data sets of two animal groups for which well-developed anatomy ontologies and/or semantic data are available: bees and fishes. Our new package provides the foundational tools to foster further advances in the field and incentivize researchers interested in working in the interface of phylogenetics, comparative methods, and ontologies.

## 2 Material and Methods

### 2.1 Implementation

*rphenoscate* is one of the two main R packages (*rphenoscape* being the other) resulting from the SCATE project (https://scate.phenoscape.org/)—Semantics for Comparative Analyses of Trait Evolution. It is tailored to facilitate comparative analyses of trait data incorporating domain knowledge from anatomy ontologies. Our package is intended as an integrative tool for comparative morphologists and systematists to work with semantic representations of organismal anatomy and/or semantically enriched phylogenetic data. The package allows working with external ontologies in OBO format but is specially integrated with the Phenoscape KB. Its sister package under development, *rphenoscape*, is tailored to work with semantic phenotypes from Phenoscape KB, including tools for quantifying the semantic similarity of phenotype descriptions and algorithms for synthesizing annotated morphological data from published studies. *rphenoscate* imports and depends on several functions from its sister package, *rphenoscape*, particularly for accessing semantic phenotypes of vertebrates available at the Phenoscape KB (https://kb.phenoscape.org/), querying absence/presence data with OntoTrace (Dececchi et al., 2015), and calculating semantic similarity metrics. It relies on *ontologyIndex* (Greene et al., 2017) for importing and working with external ontologies and *igraph* (Csardi and Nepusz, 2006) for extracting adjacency matrices and other graph manipulations. It also uses some functions from *ontoFAST* (Tarasov, 2022) to post-process semi-automatic annotations of phylogenetic character matrices with anatomy terms from the external ontologies.

### 2.2 Availability

The *rphenoscate* package requires R 3.5.0 or higher and the installation of *rphenoscape* from GitHub (https://github.com/phenoscape/rphenoscape). The current version of *rphenoscate* can be installed directly from its GitHub repository (https://github.com/uyedaj/rphenoscate). The source code from the latest stable version of the package as dated from this publication is deposited at Zenodo (XXXX).

### 2.3 Overview

The functions of *rphenoscate* comprise three main groups. The first group (G1) includes functions for: (a) assessing the dependency structure of anatomical entities based on annotations with ontology terms or semantic phenotypes available at Phenoscape KB; (b) setting up and fitting evolutionary models accounting for trait dependencies; and (c) performing stochastic character mapping using corHMM or RevBayes. The second group (G2) includes functions for: (a) assessing the relationships among anatomy ontology terms annotated to phylogenetic characters using semantic similarity metrics calculated with *rphenoscape*; and (b) visualizing the semantic and phylogenetic structure of the data. Finally, the third group (G3) includes functions for: (a) constructing phylogenetic characters based on the exclusivity classes inferred with *rphenoscape*; (b) assembling and exporting synthetic character matrices for phylogenetic analyses. A scheme of the main components in *rphenoscate* is presented in Figure 1. Detailed tutorials with examples of the different applications of *rphenoscate* are given in the Supporting Information and are also available at GitHub (https://github.com/diegosasso/rphenoscate_tutorials).

**Figure 1:**
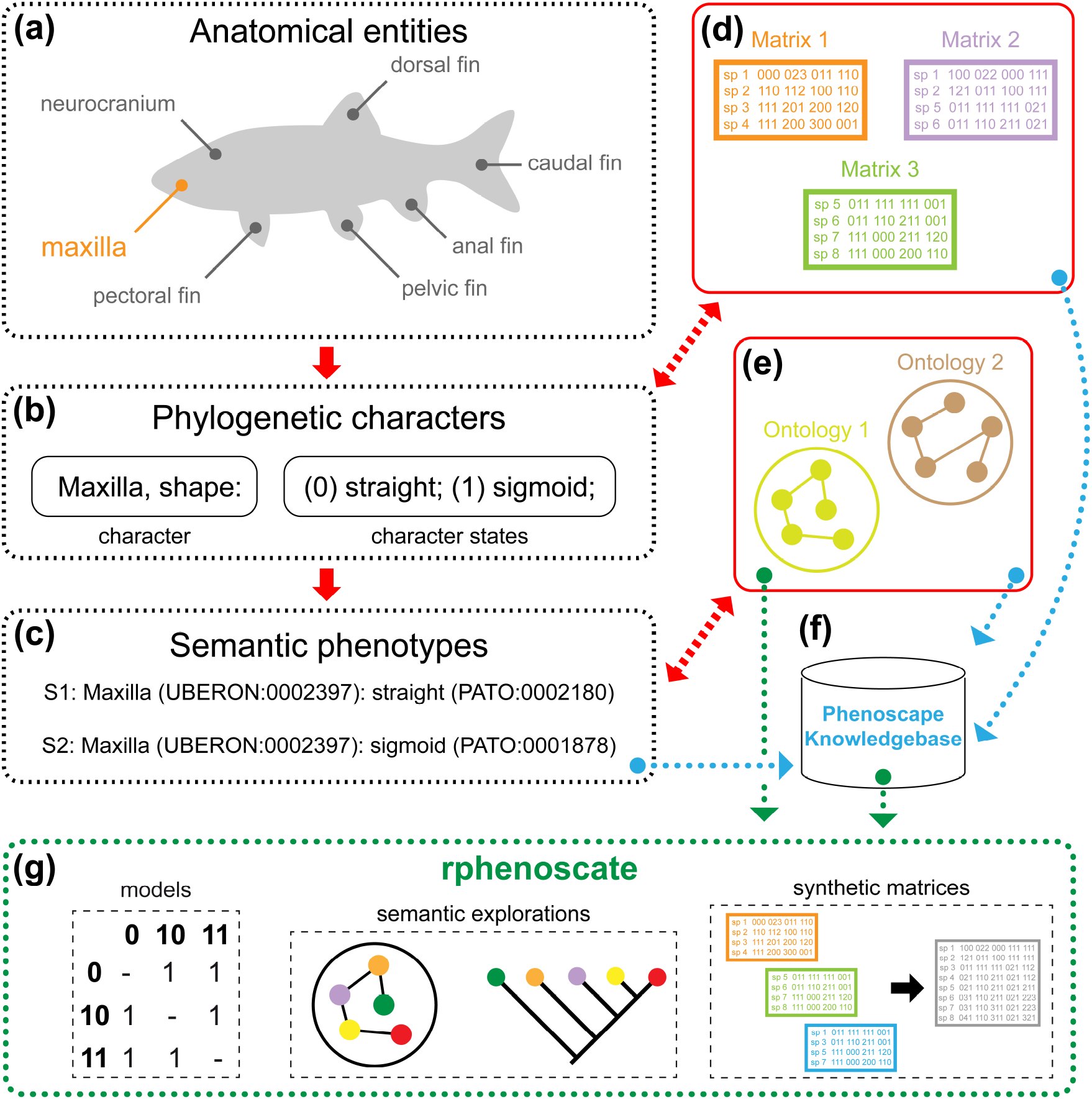
Scheme of the main concepts and components in *rphenoscate*. (a) Organismal anatomy can be conceptualized and described through anatomical entities; all of which are valuable for phylogenetic inference at a particular phylogenetic level (*e*.*g*., ‘maxilla’). (b) A systematist can thus propose a phylogenetic character formalizing the putative phylogenetic evidence; multiple phylogenetic characters evaluated for multiple taxa are usually organized in a character matrix (d). (c) An expert can further enrich the phylogenetic character with semantic information, thus proposing a semantic phenotype, by linking the anatomical entities and qualities to concepts in an anatomy ontology (e). (f) The Phenoscape Knowledgebase contains expert-curated annotations of semantic phenotypes from multiple phylogenetic studies of teleost fishes and integrates multiple ontologies (*e*.*g*., PATO, UBERON). (g) The *rphenoscate* package allows integrating knowledge from ontologies and accessing semantic phenotypes available at the Phenoscape KB (f) to automate model specification for dependent traits, perform semantic explorations of data, and assemble synthetic character matrices with the aid of the *rphenoscape*.

### 2.4 Data sets

For demonstrating the package functionality, we employed two animal groups with well-established anatomy ontologies: bees and fishes. For the bees, we employed a data set based on the matrix of corbiculate bees (Hymenoptera: Apidae; *e*.*g*., honey bees, bumble bees) from Porto and Almeida (2021) (data set 1). The original character matrix in NEXUS format was first imported in R. Then a sample of 20 phylogenetic characters referring to the anatomical entities in Table 2 was used. These characters were selected to represent anatomical entities from the head, mouthparts, and genitalia of bees, for which many anatomical dependencies can be observed (D.S.P. personal observations), making them suitable to test the package functionality. Phylogenetic character annotation used anatomy terms from the HAO ontology (Yoder et al., 2010) employing a semi-automatic pipeline implemented in *ontoFAST* (Tarasov, 2022) and new functions from *rphenoscate*.

**Table 2:** Anatomical entities and phylogenetic characters studied from the Porto and Almeida (2021) data set and corresponding terms from the HAO ontology. C1-20 denote the phylogenetic characters.

| HAO | entity | Phylogenetic character |
| --- | --- | --- |
| HAO:0000212 | clypeus | C1. Clypeus |
| HAO:0000212 | clypeus | C2. Lateral margin of ventral portion of clypeus |
| HAO:0000690 | paraocular carina | C3. Paraocular carina |
| HAO:0001454 | anterior tentorial arm | C4. Spur produced laterad from the dorsal sheet of anterior tentorial arm towards mesal margin of compound eye margin at level of antennal foramen |
| HAO:0001454 | anterior tentorial arm | C5. Lateral spur of the dorsal sheet of anterior tentorial arm |
| HAO:0001343 | posterior tentorial arm | C6. Fan-shaped sheet of posterior tentorial arm |
| HAO:0001565 | hypopharyngeal lobe | C7. Shape of hypopharyngeal lobe |
| HAO:0000456 | labrum | C8. Median tubercle or transversal ridge on distal portion of anterior surface of labrum |
| HAO:0000081 | acetabular groove | C9. Acetabular groove of mandible of female |
| HAO:0000676 | outer groove | C10. Outer groove of mandible of female |
| HAO:0000219 | condylar groove | C11. Condylar groove of mandible of female |
| HAO:0000958 | stipes | C12. Comb on an emargination at distal portion of posterior margin of stipes |
| HAO:0000958 | stipes | C13. Setae of the stipital comb |
| HAO:0000457 | lacinial lobe | C14. Shape of lacinia |
| HAO:0002149 | postarticular portion of the postmentum | C15. Mentum |
| HAO:0000686 | paraglossa | C16. Paraglossa |
| HAO:0002498 | furcula | C17. Furcula |
| HAO:0002498 | furcula | C18. Dorsal arm of furcula |
| HAO:0000707 | penisvalva | C19. Dorsal bridge of penis valves |
| HAO:0000395 | harpe | C20. Gonostylus |

For the fishes, we employed two data sets comprising skeletal characters for species in the order Characiformes (Ostariophysi). One data set was an Ontotrace (Dececchi et al., 2015) data matrix of absence/presence characters inferred for species of the family Characidae (commonly known as characids and tetras) retrieved from the Phenoscape KB (data set 2), including a search for the anatomical entities in Table 3. These entities were selected because information for them was available for many species at the Phenoscape KB and they exhibited anatomical dependencies, thus making them suitable to test the package functionality. The second data set was the matrix of anostomoid fishes (Characiformes: Anostomoidea) from Dillman et al. (2016) (data set 3). The original character matrix was retrieved from the metadata available at Phenoscape KB. This data set was selected as the benchmark of the SCATE project for evaluating the synthetic character matrix assembling functionality because the original study itself comprises a supermatrix for four families of anostomoid fishes and has semantic phenotypes available at the Phenoscape KB. For both data sets, anatomical entities were already annotated by experts (W.M.D. and P.M.M.) with anatomy terms from the UBERON ontology (Mungall et al., 2012; Haendel et al., 2014; Dahdul et al., 2018).

**Table 3:**
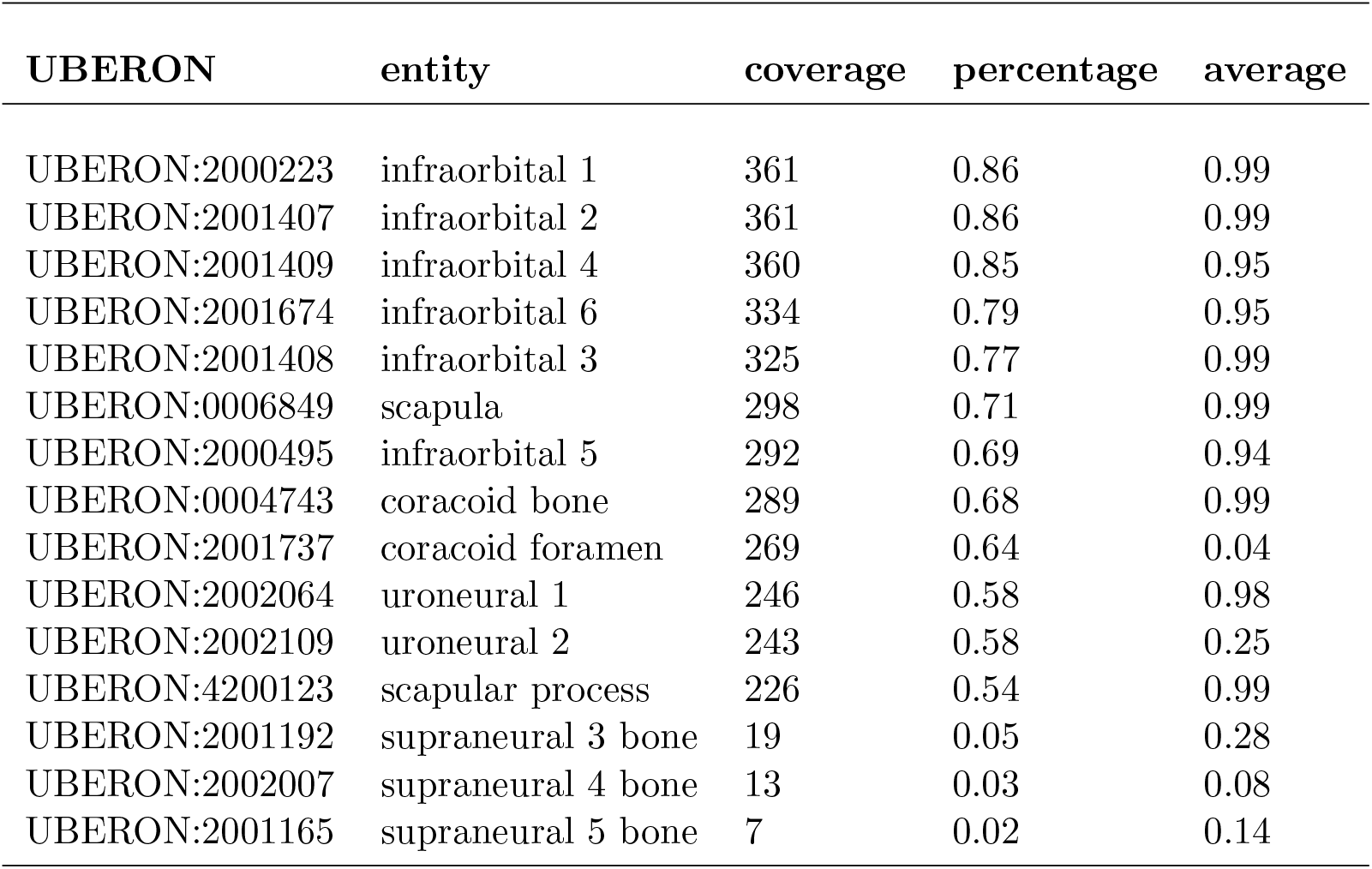
Anatomical entities studied for 420 species in the characid data set and corresponding terms from the UBERON ontology. Coverage and percentage represent respectively the number and proportion of species with absence/presence data available. Average indicates the mean presence (state 1) of anatomical entities across all species with data available.

| UBERON | entity | coverage | percentage | average |
| --- | --- | --- | --- | --- |
| UBERON:2000223 | infraorbital 1 | 361 | 0.86 | 0.99 |
| UBERON:2001407 | infraorbital 2 | 361 | 0.86 | 0.99 |
| UBERON:2001409 | infraorbital 4 | 360 | 0.85 | 0.95 |
| UBERON:2001674 | infraorbital 6 | 334 | 0.79 | 0.95 |
| UBERON:2001408 | infraorbital 3 | 325 | 0.77 | 0.99 |
| UBERON:0006849 | scapula | 298 | 0.71 | 0.99 |
| UBERON:2000495 | infraorbital 5 | 292 | 0.69 | 0.94 |
| UBERON:0004743 | coracoid bone | 289 | 0.68 | 0.99 |
| UBERON:2001737 | coracoid foramen | 269 | 0.64 | 0.04 |
| UBERON:2002064 | uroneural 1 | 246 | 0.58 | 0.98 |
| UBERON:2002109 | uroneural 2 | 243 | 0.58 | 0.25 |
| UBERON:4200123 | scapular process | 226 | 0.54 | 0.99 |
| UBERON:2001192 | supraneural 3 bone | 19 | 0.05 | 0.28 |
| UBERON:2002007 | supraneural 4 bone | 13 | 0.03 | 0.08 |
| UBERON:2001165 | supraneural 5 bone | 7 | 0.02 | 0.14 |

### 2.5 Package showcase

For showcasing the package, we consider three study cases, one for each data set presented above. In the first case (hereafter BEES), a researcher wants to reconstruct the evolutionary history of several traits and understand how they relate to each other in bee anatomy (data set 1). For example, do traits from different anatomical regions evolve similarly? How are the anatomical entities represented by such traits related to each other? The researcher needs first to account for possible dependencies among anatomical entities in the evolutionary models (*i*.*e*., biologically realistic models) before reconstructing the trait histories using stochastic character mapping. Then, the researcher needs to employ some tool to visualize the semantic patterns across anatomical entities in the data.

In the second case (hereafter CHARA), a researcher has access to the Phenoscape KB and wants to retrieve all information available for absence/presence of bones in characid fishes (data set 2). The researcher wants then to reconstruct the evolutionary history of these traits to answer a particular question. Do bones from particular body regions get lost more frequently than others in this particular group of fishes? For that, this researcher also needs to account for possible dependencies among anatomical entities when reconstructing character histories and employ tools to investigate the association between the semantic and phylogenetic patterns of the data.

In the third case (hereafter ANOST), a researcher also has access to the Phenoscape KB but this time wants to retrieve all information available for semantic phenotypes in anostomoid fishes. The researcher wants then to use this information to infer a phylogenetic tree. For that, the researcher needs some tools for getting the semantic phenotypes (task 1), converting them to phylogenetic characters (task 2), and assembling them in a synthetic character matrix (task 3). However, how can this researcher be assured that a synthetic character matrix obtained as such actually contains phylogenetic information? To answer this question, a benchmark is necessary, thus the matrix of anostomoid fishes from Dillman et al. (2016) (data set 3) was used.

### 2.6 Assessment analyses

BEES and CHARA.—Stochastic character mapping was used to reconstruct trait evolution using corHMM (Boyko and Beaulieu, 2021). For BEES, reconstructions used an ultrametric tree modified from Porto and Almeida (2021) using *phytools* (Revell, 2012). Note that the transformation was done only for demonstrative purposes and a proper dating method was not employed. For CHARA, reconstructions used a dated phylogeny obtained from *fishtree* (Chang et al., 2019). In both cases, for the exploration of the semantic patterns of the data, clustering dendrograms for the anatomy ontology terms (‘trait trees’) were constructed using the Jaccard semantic similarity metric calculated using functions from *rphenoscape*.

ANOST.—Assessment of phylogenetic information was performed by comparing the original data set from Dillman et al. (2016) to the synthetic character matrix obtained from semantic phenotypes of the same study using functions from *rphenoscape* (tasks1 and 2) and *rphenoscate* (task 3). Comparisons were made for both character matrices and for the posterior distributions of trees inferred from them. Character matrices were compared by calculating the cladistic information content (*sensu* Steel and Penny, 2005) using functions from the package *TreeTools* (Smith, 2019). Posterior distributions were compared by calculating the generalized Robinson-Foulds (RF) distances (Smith, 2020a) in reference to the majority-rule (MJ) consensuses of both analyses using functions from the package *TreeDist* (Smith, 2020b). The generalized RF distance is a metric of dissimilarity between pairs of trees based on the information content (in bits) of shared splits (Smith, 2020b). In short, posterior distributions of tree topologies were sampled through Bayesian inferences for both character matrices. Then generalized RF distances were calculated in reference to the MJ consensus of the original and inferred synthetic matrices, thus resulting in four distributions of RF distances: distribution from the (i) original matrix vs. original MJ consensus; (ii) inferred synthetic matrix vs. original MJ consensus; (iii) inferred synthetic matrix vs. inferred synthetic MJ consensus; and (iv) original matrix vs. inferred synthetic MJ consensus. A broad overlap between (i) and (ii) and between (iii) and (iv) can then serve as a proxy to assess whether the Bayesian phylogenetic analyses result in similar posterior distributions of trees and thus whether character matrices have similar phylogenetic information. Bayesian inferences were performed in MrBayes (Ronquist et al., 2012) with MCMC settings as indicated in the Supporting Information available online.

Finally, to give an example based on the original intent of the researcher in this study case, an additional search was performed retrieving all semantic phenotypes available at Phenoscape KB for fishes in Characidae. This family was selected—instead of the superfamily Anostomoidea—to reduce computational effort and facilitate downstream analyses (for demonstrative purposes only), but still, show an example of a relatively large data set retrieved from Phenoscape KB. The data set was then used to build a synthetic character matrix assembling data from multiple phylogenetic studies (see also Dececchi et al., 2015).

## 3 Results

### Automated construction of structured Markov models for dependent traits and exploration of semantic patterns of morphological data

BEES.—The sample of 20 phylogenetic characters from Porto and Almeida (2021) contained 16 anatomical entities (Table 2). In those cases where multiple characters refer to the same anatomical entity, *rphenoscate* automatically detected those characters and set up appropriate evolutionary models, either a standard structured Markov model (SMM-ind) if no ontological dependencies were found; an embedded dependency quality type Markov model (ED-ql) if dependencies based on property instantiation were found (*sensu* Vogt, 2018a); or an embedded dependency absence-presence type Markov model (ED-ap) if dependencies based on parthood relations were found (*sensu* Vogt, 2018a; for additional discussions on types of dependencies and models see Tarasov, 2019, 2022; Vogt, 2018a). Otherwise, different models were automatically assigned to single non-dependent characters based on the number of observed states (Figure 2). For example, amalgamated characters of the ‘posterior tentorial arm’ were assigned an Mk model with 2 states; the ‘anterior tentorial arm’, an ED-ql model with 3 states; the ‘furcula’, an ED-ql model with 6 states; and the ‘hypopharyngeal lobe’, an Mk model with 7 states. Samples of the stochastic maps from these examples are shown in Figure 3a.

**Figure 2:**
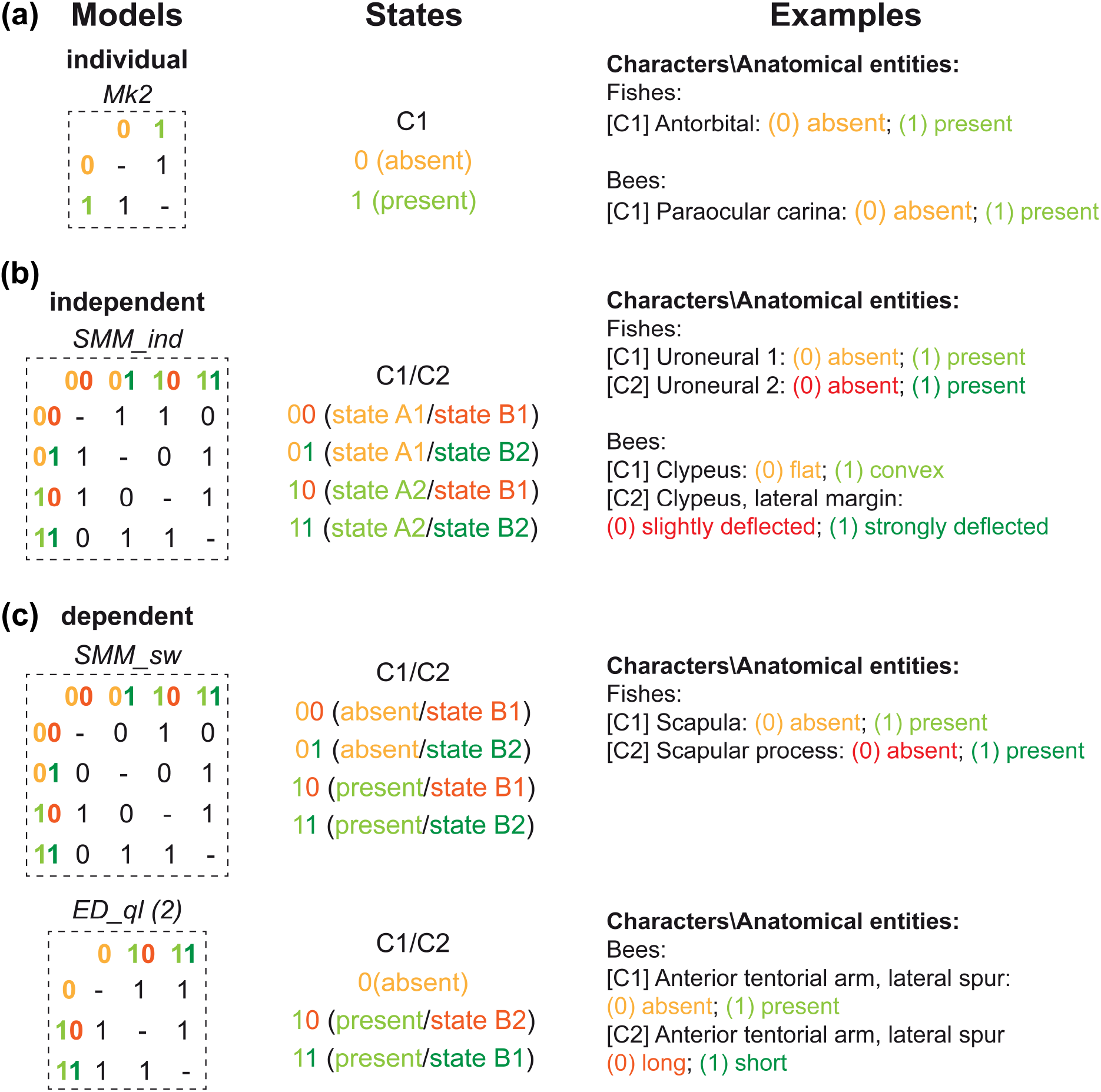
Types of models automatically set-up by *rphenoscate*. (a) Standard Markov models with variable number of states for individual characters (Mk), in this case, a binary character. (b) Structured Markov models for groups of independent characters (SMM-ind), in this case, a pair of binary characters. (c) Two types of models that account for character dependencies: Structured Markov models of the switch-on type (SMM-sw) and embedded dependency Markov models of the quality type (ED-ql). In both cases, the example is for a pair of binary characters. Note that SMM-sw and ED-ql treat absences differently (state 0); as two combinations of hidden states (only one observable) in the former and only one observable state in the latter. C1 and C2 indicate characters 1 and 2 respectively. Color codes are used to facilitate character state visualization for characters.

**Figure 3:**
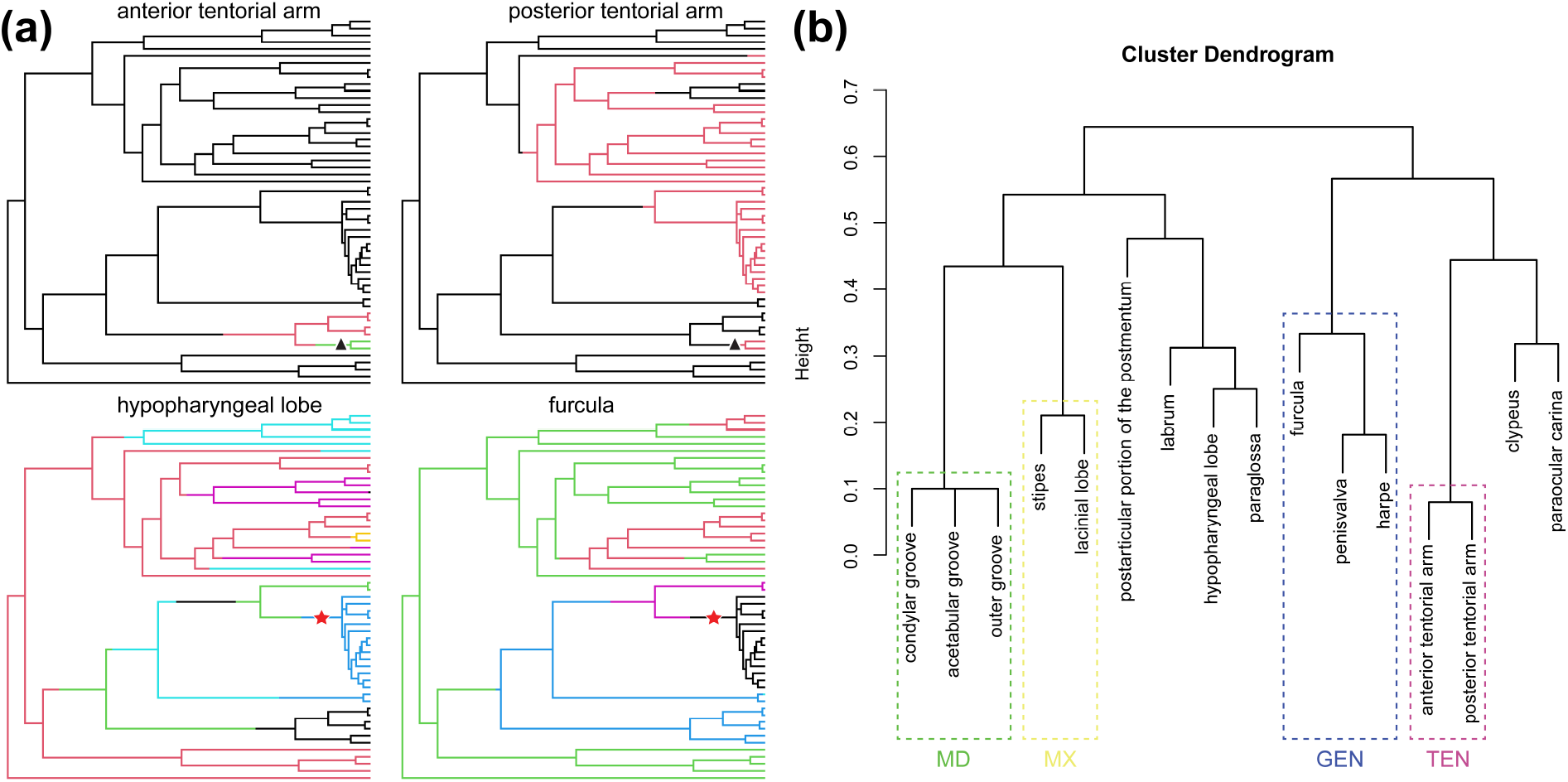
Exploration of the bee data set of Porto and Almeida (2021). (a) Sample of stochastic character maps obtained from four different anatomical entities. Branches in different colors indicate different ancestral character states. The red stars and black triangles indicate some clades with congruent patterns of reconstructed character states. (b) Clustering dendrogram showing the relationships among HAO terms referring to the anatomical entities of this data set based on the Jaccard semantic similarity. Dashed boxes indicate some clusters based on parthood relations known for the Hymenoptera anatomy. Abbreviations: GEN, genitalia; MD, mandible; MX, maxilla; TEN, tentorium.

In this study case, the researcher was interested in reconstructing the evolutionary history of multiple traits and understanding their relationships in the bee anatomy. After accounting for the ontological dependencies among anatomical entities in the evolutionary models, the researcher can observe that reconstructed trait histories show some character states co-occurring in the phylogeny, for example, those in the clades indicated with stars and triangles (Figure 3a). When exploring the semantic patterns of the data, the relationships among the ontology term annotations indicate that some anatomical entities are part of the same anatomical regions of the bee anatomy (*e*.*g*., ‘anterior tentorial arm’ and ‘posterior tentorial arm’ are *part of* ‘tentorium’; Figure 3b, TEN, purple dashed box) whereas others not (*e*.*g*., ‘hypopharyngeal lobe’ and ‘furcula’). Most clusters based on semantic similarity, in this case, actually correspond to anatomically related entities of the bee anatomy, as indicated by parthood relationships to parent terms in the HAO ontology. For example, clusters with anatomical entities that are *part of* ‘mandible’, ‘maxilla’, ‘genitalia’, and ‘tentorium’ were recovered (dashed boxes in Figure 3b; MD, MX, GEN, and TEN respectively). Therefore, clustering anatomical entities based on semantic similarity metrics calculated for their ontology term annotations can be used by this researcher to further investigate if such clusters reflect shared parameters in the evolutionary models of traits linked to these anatomical entities, for example, evolutionary rates or transition biases.

CHARA.—The data set retrieved from the Phenoscape KB contained 420 species with absence/presence data available for at least one of the anatomical entities listed in Table 3. From these, 146 species were also available in the tree obtained from *fishtree*. Data coverage, defined as the number of species for which absence/presence was asserted or can be inferred by the Phenoscape KB reasoner, ranges from 361 (86%) to 7 (2%) across all taxa (Table 3). The average presence of anatomical entities across species with data available ranges from 0.99 for ‘infraorbital 1’ and ‘infraorbital 2’ to 0.14 for ‘supraneural 5 bone’, with lower values indicating entities commonly absent (*e*.*g*., ‘coracoid foramen’, ‘uroneural 2’, ‘supraneural bone’, ‘supraneural 4 bone’). From the anatomical entities in Table 3, ontological relationships were detected between the pairs ‘scapula’ and ‘scapular process’, and ‘coracoid bone’ and ‘coracoid foramen’, thus appropriate structured Markov models were automatically set up by *rphenoscate*. In this case, the model used to account for trait dependencies was the SMM-sw, as discussed in Tarasov (2019, 2022), as shown in Figure 2. Samples of stochastic maps for some of the anatomical entities, including the two above pairs of dependent ones, are shown in Figure 4.

**Figure 4:**
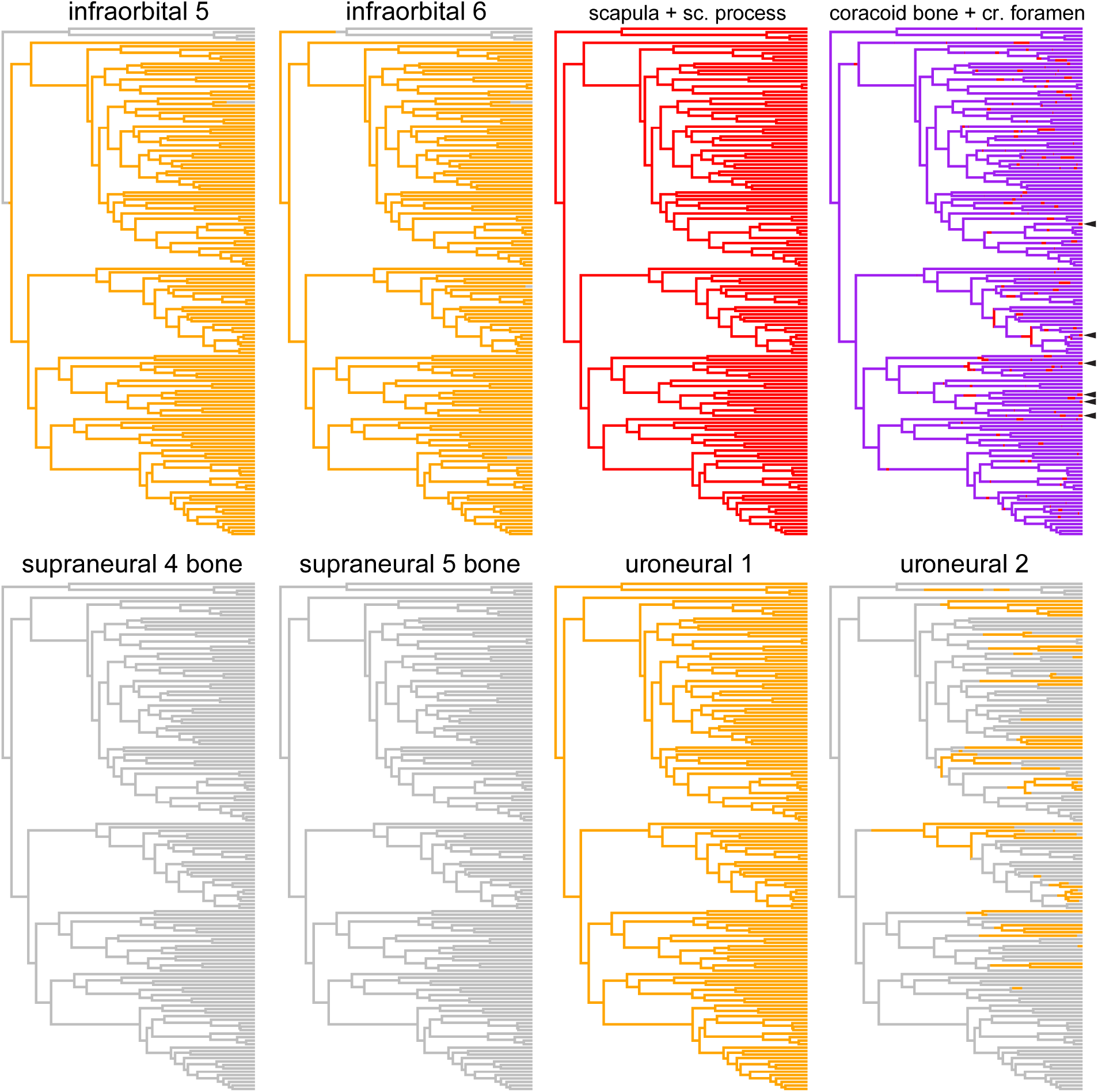
Sample of stochastic character maps obtained from ten different anatomical entities of the Characidae data set. Branches in orange indicate inferred presence of the respective anatomical entity and those in grey indicate absence; for pairs of entities, red color indicates the presence of both, as indicated with arrowheads for the pair ‘coracoid bone + coracoid foramen’ and purple indicates the presence of the first entity but the absence of the second.

As observed for ‘supraneural 4 bone’, ‘supraneural 5 bone’, and ‘uroneural 1’, for example, stochastic character maps reconstructed no transitions at all, possibly due to many taxa being coded as polymorphic (*i*.*e*., states 0 and 1 or 1 and 0) or ‘?’ (missing) and/or due to low data coverage, as is observed in ‘supraneural 4 bone’ and ‘supraneural 5 bone’. In the case of the combined character ‘coracoid bone + coracoid foramen’, all instances of presence of ‘coracoid foramen’ seem to be correctly inferred in branches where ‘coracoid bone’ was also present, as indicated with the arrowheads in Figure 4.

In this second study case, the researcher was interested in understanding the history of loss of bones in characid fishes. By inspecting the stochastic character maps (4), the researcher can observe that some bones were reconstructed as absent (*e*.*g*., ‘supraneural 4 bone’ and ‘supraneural 5 bone’) or present for all species (*e*.*g*., ‘uroneural 1’), possibly due to the issues mentioned above. Some other bones were lost multiple times in several species (*e*.*g*., ‘uroneural 2’) whereas others were lost a few times but seem to be correlated (*e*.*g*., ‘infraorbital 5’ and ‘infraorbital 6’). More complex cases can be observed for the combined characters. For example, for ‘scapula + scapular process’, ‘scapula’ and ‘scapular process’ are present in all species, whereas for ‘coracoid bone + coracoid foramen’, ‘coracoid bone’ is present in all species, but ‘coracoid foramen’ can be absent or present (4, arrowheads).

However, the researcher can learn more about the losses of bones in characid fishes by also investigating the semantic patterns of the data with some tools from *rphenoscate*. For example, in Figure 5, the tree shown to the left is the species phylogeny obtained from the *fishtree* package; the clustering dendrogram at the top right shows the relationships among the anatomical entities from Table 3; and the heatmap indicates absence/presence of the bones. In this case, some phylogenetic patterns of the data set can be easily identified, such as the absence of ‘infraorbital 5’ and ‘infraorbital 6’ supporting the clade indicated with a red dashed-box in the phylogenetic tree of Figure 5. Additionally, a clear pattern in this data set is that information-poor anatomical entities—empty cells in the heatmap—are not randomly distributed; rather they are predominantly semantically related entities: all bones from the supraorbital series (Figure 5: clustering dendrogram, star). This might prompt the researcher to further investigate if this lack of information is simply due to a poorly studied anatomical structure in this group of fishes or if there are underlying biological causes.

**Figure 5:**
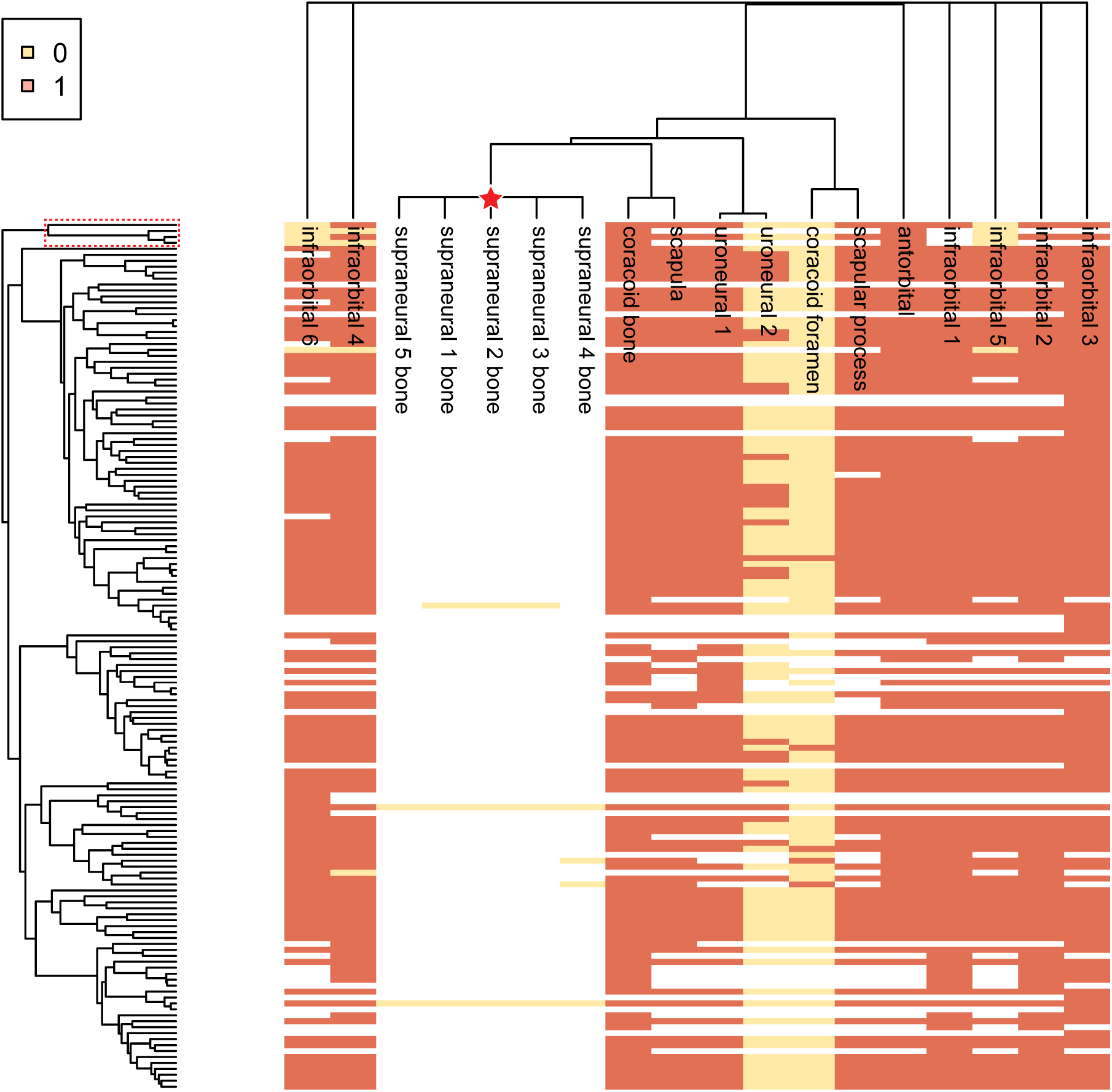
Visualization of phylogenetic and semantic patterns of the Characidae data set. The tree to the left is the dated species phylogeny obtained from the *fishtree* package. The clustering dendrogram at the top shows the relationships among UBERON terms referring to the anatomical entities of this data set based on the Jaccard semantic similarity. The heatmap shows absence (state 0, yellow) or presence (state 1, orange) for each anatomical entity in each species; empty cells indicate the absence of information. The dashed box at the top of the phylogeny indicates a clade of fishes supported by the absence of the bones ‘infraorbital 5’ and ‘infraorbital 6’. The red star in the dendrogram indicates a cluster of related anatomical entities with a lack of information for this particular group of fishes.

### Synthetic character matrices maintain phylogenetic information from manually-curated matrices

ANOST.—The ability to synthesize data from different studies with characters of varying types presents a major challenge to data reuse, expansion, and synthesis (Dececchi et al., 2015). In this third study case, the researcher was interested in retrieving all semantic phenotypes for anostomoid fishes from the Phenoscape KB, building a character matrix, and inferring a phylogeny. However, this task requires assessing the phylogenetic utility of this synthetic character matrix. For that, the researcher evaluated whether the use of character data represented as ontology-annotated phenotypic statements (‘semantic phenotypes’) and subsequent construction of synthetic character matrices from these phenotypes resulted in any loss of phylogenetic information. The researcher achieved this by using *rphenoscate* and *rphenoscape* to compare the semantic phenotypes obtained from the Phenoscape KB to the original expert-curated matrix from Dillman et al. (2016).

The original data set from Dillman et al. (2016) contained 463 phylogenetic characters and 173 taxa. With *rphenoscate*, it was possible to recover and cluster semantic phenotypes referring to the original data set resulting in a synthetic matrix with 422 characters. When assessing the phylogenetic information of both data sets, the cladistic information content (*sensu* Steel and Penny, 2005) for characters in the original and synthetic matrices are almost identical (Figure 6a) indicating the conservation of potential phylogenetic information (Porto et al., 2022). When comparing the majority-rule consensus trees inferred from both matrices (Figure 6b) or their posterior distributions (Figure 6c-d), trees are almost identical and distributions mostly overlap, demonstrating that the phylogenetic information of the original data set was retained in the synthetic matrix inferred with *rphenoscate*.

**Figure 6:**
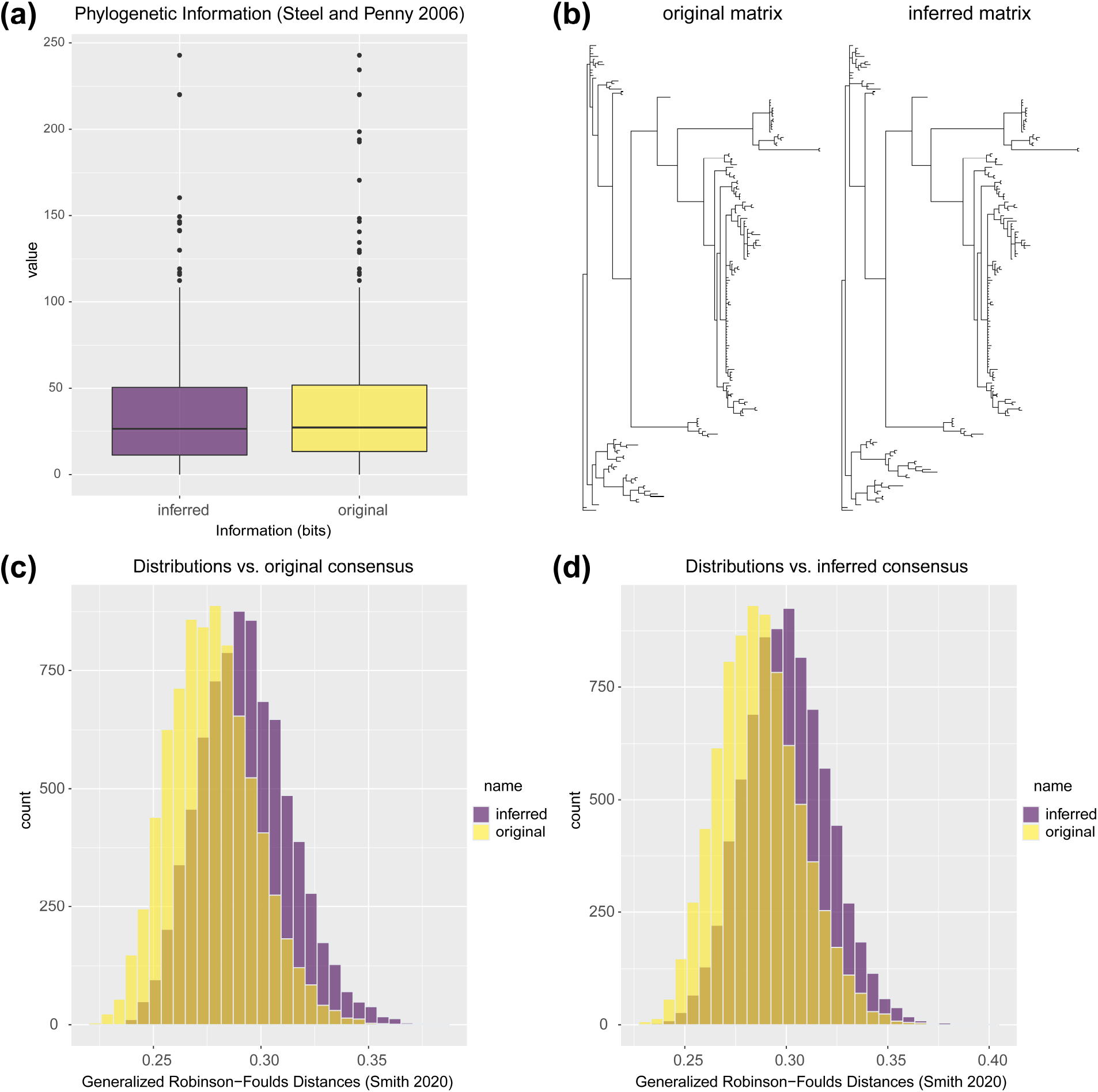
Assessments of the phylogenetic information of the original and inferred synthetic anostomoid data sets from Dillman et al. (2016). (a) Boxplots of cladistic information content (*sensu* Steel and Penny, 2005) for phylogenetic characters in both data sets. (b) Majority-rule consensus trees inferred from Bayesian analyses of both data sets. (c) Distribution of Generalized Robinson-Foulds distances for trees in the posterior obtained from the original and inferred synthetic data sets compared to the majority-rule consensus tree of the original data set. (d) Same as (c) but compared to the majority-rule consensus tree of the inferred data set.

As for the additional search on the Phenoscape KB, the synthetic matrix inferred from semantic phenotypes of Characidae contained 524 species and 739 phylogenetic characters. From all species, around 45% have data available for at least a quarter of the phylogenetic characters. From all phylogenetic characters, at least 37% have data available for at least a quarter of the species. Overall data coverage—character state information available—is around 20% for the entire matrix (Figure 7). From all phylogenetic characters, around 20% are phylogenetically non-informative (*i*.*e*., non-variable for the taxa considered).

**Figure 7:**
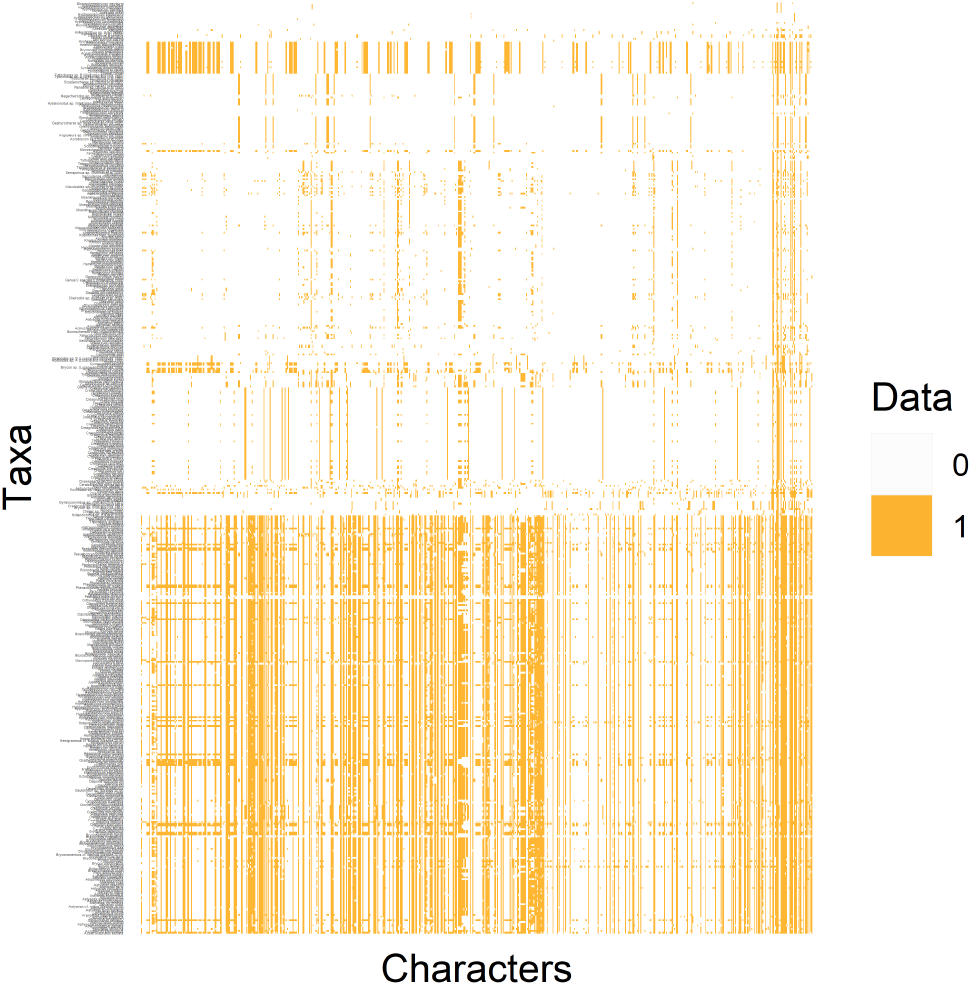
Heatmap representing the synthetic character matrix obtained from all semantic phenotypes of Characidae available at Phenoscape KB. Filled cells (state 1, orange color) indicate information available for a given taxon, irrespective of the actual character state.

A complete work-through of all the analyses of the three study cases is given in the tutorials in the Supporting Information online and also available on GitHub (https://github.com/diegosasso/rphenoscate_tutorials).

## 4 Discussion

### 4.1 Studying complex traits

One of the main challenges of studying morphological evolution is modeling complex traits—sets of related traits often exhibiting multiple levels of dependencies or correlations (*e*.*g*., Tarasov, 2022: Fig. 1D). We have demonstrated that morphological knowledge expressed in anatomy ontologies can be employed for automatically setting up models for ontologically dependent traits. Biologically realistic models for morphology—*e*.*g*., accounting for ontological dependencies or correlations among characters— can be used for studying complex traits, for example, in the context of understanding adaptations to particular environments (Tribble et al., 2022); trait-dependent diversification (O’Meara et al., 2016); or integration/modularity among anatomical structures (Billet and Bardin, 2019).

### 4.2 What can be learned from the three study cases?

Trait evolution and semantic patterns.—In this work, we have shown the application of ontology-informed evolutionary models for morphological traits in the context of stochastic character mapping with two data sets, bees (Figure 3) and characid fishes (Figure 4), annotated with terms from the HAO and UBERON ontologies, respectively. We then demonstrated how *rphenoscate* can help researchers to investigate trait evolution and address simple evolutionary questions by assessing semantic patterns in morphological data.

In the study-case BEES, after accounting for ontological (anatomical) dependencies among traits, the researcher learned that some character states are still reconstructed in similar branches of the tree (stars and triangles in Figure 3a). Although this pattern is congruent with a scenario of biological dependency between traits, the limited size of the data set—only one instance of co-occurring states— precludes any assertive interpretation. Another possibility is that some traits from structurally related anatomical regions might be evolving similarly due to shared gene regulatory and developmental machinery (Wagner Gunter and Altenberg, 1996; Wagner and Stadler, 2003; Mabee, 2006). By investigating the semantic patterns of ontology annotations to phylogenetic characters in this data set, the researcher learned that some traits with congruent character-state reconstructions (triangles in Figure 3a) represent related anatomical entities—*i*.*e*., that are *part of* the same anatomical region (Figure 3b, TEN, purple dashed box). Indeed, this might be an indicator that some traits from a given anatomical region evolve similarly. However, in the context of phylogenetic inference, it has been demonstrated that the evolution of morphological characters does not necessarily follow anatomical partitions (Tarasov and Genier, 2015; Casali et al., 2022) or is often incongruent across them (Porto et al., 2021, 2022), thus prompting the researcher to further investigate for alternative causal explanations.

In the study-case CHARA, the researcher learned that some bones representing structurally related anatomical entities might be evolving independently (*e*.*g*., ‘uroneural 1’ and ‘uroneural 2’) whereas others not (*e*.*g*., infraorbital bones) (Figure 4). They could also observe that anatomical entities commonly lost in characid fishes include both structurally related (*e*.*g*., ‘supraneural 3 bone’, ‘supraneural 4 bone’, and ‘supraneural 5 bone’) and unrelated entities (*e*.*g*., ‘coracoid foramen’ and ‘uroneural 2’) (Figure 5). Furthermore, the loss of some structurally related entities (*e*.*g*., infraorbital bones) seems to be phylogenetically informative for some groups of fishes (Figure 5, red dashed box). After this initial exploration using *rphenoscate*, the researcher can then investigate the observed phylogenetic and semantic patterns of the data to ask further questions. For example, why are these particular bones absent altogether in some groups of fish? Are they associated with (*develops from*) the same developmental module?

Synthetic character matrices.—Finally, in the study case ANOST, the researcher was able to obtain a synthetic character matrix from semantic phenotypes and learned that the phylogenetic information inferred from this matrix is indeed comparable to that inferred from the original manually-curated matrix (Figure 6). This result is crucial since the main interest of most systematists in assembling character matrices is to infer the phylogeny of a given group based on the available anatomical evidence. Perhaps more importantly, it was demonstrated that it is also possible to construct synthetic character matrices from semantic phenotypes of multiple different studies, as obtained for characid fishes (Figure 7). This opens up opportunities for ‘phenomic-scale’ studies with synthetic matrices (*e*.*g*., Dececchi et al., 2015) exploring all the semantic phenotypes of teleost fishes available at Phenoscape KB and provides a model for future knowledgebases focused on other groups of organisms.

### 4.3 Current limitations

Although *rphenoscate* offers some tools for working with external ontologies (*i*.*e*., other than UBERON) and NEXUS files associated with ontology annotations, other tools are specifically for working with the semantic phenotypes from the Phenoscape KB in synergy with *rphenoscape*. Furthermore, a major limitation in both cases—external character matrices or Phenoscape KB data—is that annotation of phylogenetic characters with ontology terms has to be done manually. In the case of external ontologies, semi-automatic annotations can be performed using *ontoFAST* and post-processing with *rphenoscate*, but those are limited to the anatomy terms only (*i*.*e*., thus not including quality terms describing character states) and the final decision still requires expert judgment. One additional limitation is the number of models currently implemented to account for dependencies using ontology information (ED-ap, ED-ql, and SMM) and the automatic setting up option being restricted to only linear chains of dependencies and a few hierarchical levels (*e*.*g*., entity A *depends on* entity B; or entity A *depends on* entity B *depends on* entity C).

### 4.4 Semantic phenotypes and new approaches to morphological data

Ontologies can provide a new framework for representing and studying organismal anatomy. As suggested in Vogt (2018a,b), alternative formalizations of morphological data, other than free-text descriptions in natural language or standard character matrices, offer several new opportunities but also challenges. Some advantages of working with semantically-enriched representations of morphological information (*e*.*g*., Balhoff et al., 2010, 2014; Dececchi et al., 2015; Deans et al., 2015; Thessen et al., 2020; Stefen et al., 2022) include the possibility of automatically assembling synthetic character matrices for phylogenetic inference, as demonstrated here; integrating anatomical information at phenomic scale across databases and domains of knowledge; and developing graph-based phylogenetic algorithms for comparative analyses (Ramírez and Michalik, 2014; Vogt, 2018a,b). *rphenoscate* represents an important step in these directions.

In a broader context, working with ontologies and semantic representations of organismal anatomy have utilities and advantages beyond the few ones presented here. It is a fundamental and necessary step for fully exploiting morphological data in this new era of ‘Phenomics’ (Braun et al., 2018). It allows data from different sources and domains of knowledge to be easily integrated and summarized, making it easily findable, accessible, interoperable, and reusable by humans and machines, thus compliant with the FAIR principles in data science (Wilkinson et al., 2016). In this study, we showed that semantic phenotypes can be automatically converted into reasonable synthetic character matrices for downstream analysis. Thus, computer-assisted phenomic-scale research can be made possible in evolutionary biology. We hope that our new package will offer some useful tools in this direction encouraging interested researchers and prompting advances in the fields of comparative morphology, phylogenetics, and ontologies.

## Acknowledgments

This work received funding from the National Science Foundation (NSF 1661516 to J.C.U., NSF 1661529 to W.M.D. and P.M.M., NSF 1661456 to H.L., NSF 1661356 to T.J.V. and J.P.B.) and the Academy of Finland (339576 to S.T.).

## Author’s Contributions

D.S.P., S.T., and J.U. conceived the package. D.S.P., S.T., H.L., C.C, and J.U. designed the package, tested the package, and wrote the documentation. All authors wrote the first draft of the manuscript and revised the final version of the paper.

## Data Availability Statement

The code of *rphenoscate*, tutorials and data sets are available on GitHub at https://github.com/uyedaj/rphenoscate and https://github.com/diegosasso/rphenoscate_tutorials, and Zenodo at XXXX.

